# Hnrnpul1 controls transcription, splicing, and modulates skeletal and limb development in vivo

**DOI:** 10.1101/2020.02.04.934257

**Authors:** Danielle L Blackwell, Sherri D Fraser, Oana Caluseriu, Claudia Vivori, Amanda V Tyndall, Ryan E Lamont, Jillian S Parboosingh, A Micheil Innes, François P Bernier, Sarah J Childs

## Abstract

Mutations in RNA binding proteins can lead to pleiotropic phenotypes including craniofacial, skeletal, limb and neurological symptoms. Heterogeneous Nuclear Ribonucleoproteins (hnRNPs) are involved in nucleic acid binding, transcription and splicing through direct binding to DNA and RNA, or through interaction with other proteins in the spliceosome. We show a developmental role for Hnrnpul1 in zebrafish, resulting in reduced craniofacial tendon length, severe adult-onset scoliosis and reduced fin size. We demonstrate a role of Hnrnpul1 in alternative splicing and transcriptional regulation using RNA sequencing. Given its cross-species conservation and role in splicing it would not be surprising if it had a role in human development but the developmental role of this gene in humans has not been explored. Whole exome sequencing detected a frameshift variant in *HNRNPUL1* in two siblings with congenital limb malformations which remain variants of unknown significance. Zebrafish Hnrnpul1 mutants suggest an important developmental role of hnRNPUL1 and provide motivation for exploring potential conservation of ancient regulatory circuits involving hnRNPUL1 in human development.

## Introduction

Mutations in proteins involved in alternative splicing (AS) lead to spliceosomopathies in humans. Despite being expressed ubiquitously, mutations in core and alternative splicing factors can result in tissue-restricted, cell-type specific phenotypes including craniofacial, limb, skeletal and neurological syndromes (Lehalle *et al.*, 2015). Tissue-specificity can occur because of the sensitivity of individual tissues during embryonic development to AS, such as sensitivity of the neural crest in the case of craniofacial anomalies (Lehalle *et al.*, 2015). AS is the process of producing multiple different mRNA transcripts and protein isoforms through the differential selection of splice sites within a single pre-mRNA molecule. AS of pre-mRNA is carried out by the spliceosome, which is a complex of small nuclear RNAs and proteins. AS events mainly include exon skipping, intron retention and alternative 5’ or 3’ splice sites. For example, mutations in *TXNL4A* (Wieczorek *et al.*, 2014), *EIF4A3* (Favaro *et al.*, 2014), *EFTUD2* (Lines *et al.*, 2012), *SF3B4* (Bernier *et al.*, 2012) and *SNRPB* (Lynch *et al.*, 2014) cause human spliceosomopathies (Lehalle *et al.*, 2015).

Members of the heterogeneous nuclear ribonucleoprotein (hnRNP) family are involved in nucleic acid binding, splicing and transcription. They are present in the spliceosome and contribute directly and indirectly to the processing of pre-mRNA into mature mRNA, with nearly all hnRNP proteins having RNA-binding motifs (Dreyfuss *et al.*, 1993; Geuens *et al.*, 2016). Pathogenic variants associated with human disease have been discovered in hnRNP family members *HNRNPK* (Au *et al.*, 2015), *HNRNPU* (Poot, 2019) and *HNRNPDL* (Batlle *et al.*, 2020). In particular, the hnRNPU family often act as repressors of mRNA splicing (Matlin *et al.*, 2005). hnRNPU proteins are involved in DNA repair (Hegde *et al.*, 2012; Polo *et al.*, 2012) and U2 snRNP maturation (Xiao *et al.*, 2012). hnRNPUL1 (also known as E1B-AP5), is also a transcriptional repressor (Kzhyshkowska *et al.*, 2003).

As little is known about hnRNPUL1 function *in vivo*, we studied its developmental role in the zebrafish model. Zebrafish is an ideal model for this study due to conservation of developmental processes and genetic networks with human, coupled with rapid development. We identify a multi-tissue phenotype involving fin, craniofacial and skeletal abnormalities including scoliosis. Through RNA sequencing and alternative splicing analysis, we show that alterative splicing events are disrupted in zebrafish *hnrnpul1* mutants. We detect a homozygous frameshift variant in Heterogeneous Nuclear Ribonucleoprotein U Like 1 (HNRNPUL1) gene in two siblings with craniofacial and limb anomalies in humans, raising the possibility that HNRNPUL1 is critical to vertebrate development.

## Results

### Generation of double homozygous zebrafish *hnrnpul1* and *hnrnpul1l* mutants

hnRNPUL1 has been studied in cell lines, however there are no data on its role at the organismal level, nor during animal development. Therefore, we investigated its role in the zebrafish because of ease of genetic analysis. Due to the genome duplication in the teleost lineage, there are two closely related *HNRNPUL1* orthologues in zebrafish, *hnrnpul1* and *hnrnpul1l* and therefore we used a double knockout strategy (hereafter referred to as *hnrnpul1/1l* mutants. Human HNRNPUL1 protein sequence shows 65% similarity with zebrafish Hnrnpul1 (chromosome 18) and 67% with Hnrnpul1l (chromosome 5) respectively. The DNA binding (SAP) and protein-protein interaction (SPRY) domains show higher conservation (76% and 77% similarity for Hnrnpul1 and 83% and 78% similarity for Hnrnpul1l respectively; Fig. 1A). As we had identified two siblings with craniofacial and limb malformations with a homozygous frameshift variant of unknown significance (VUS) in HNRNPUL1 (NM_007040.5:c.1673dup, p. (Glu560Argfs*17)) we targeted CRISPR guides to make loss of function alleles in zebrafish with mutations near the human mutation site. To ensure loss of function mutation, a homology-directed repair ‘stop cassette’ with stop codons in three reading frames was included in the CRISPR-Cas9 injections. *hnrnpul1l*^Ca52^ and *hnrnpul1*^Ca53^ and *hnrnpul1*^Ca54^ alleles were isolated with mutations at similar locations to the human mutation in the DNA (Fig. S1) and protein (Fig. 1B). *hnrnpul1l*^Ca52^ has a 106-nucleotide insertion, resulting in a frame shift mutation and premature stop codon (Fig. S1B). This is predicted to produce a nonsense protein mutation resulting in truncation 3 amino acids after the mutation (Fig. 1B). *hnrnpul1*^Ca53^ has a 35-nucleotide insertion and *hnrnpul1*^Ca54^ has a 63-nucleotide insertion (Fig. S1C, D). Both mutations result in a frame shift and premature stop codon resulting in truncation 6 amino acids after the mutation for allele Ca53 and 11 amino acids for Ca54 (Fig. 1B). All three alleles are predicted to result in truncated protein. No phenotypic differences in the *hnrnpul1* Ca53 and Ca54 alleles were noted.

**Figure 1.**
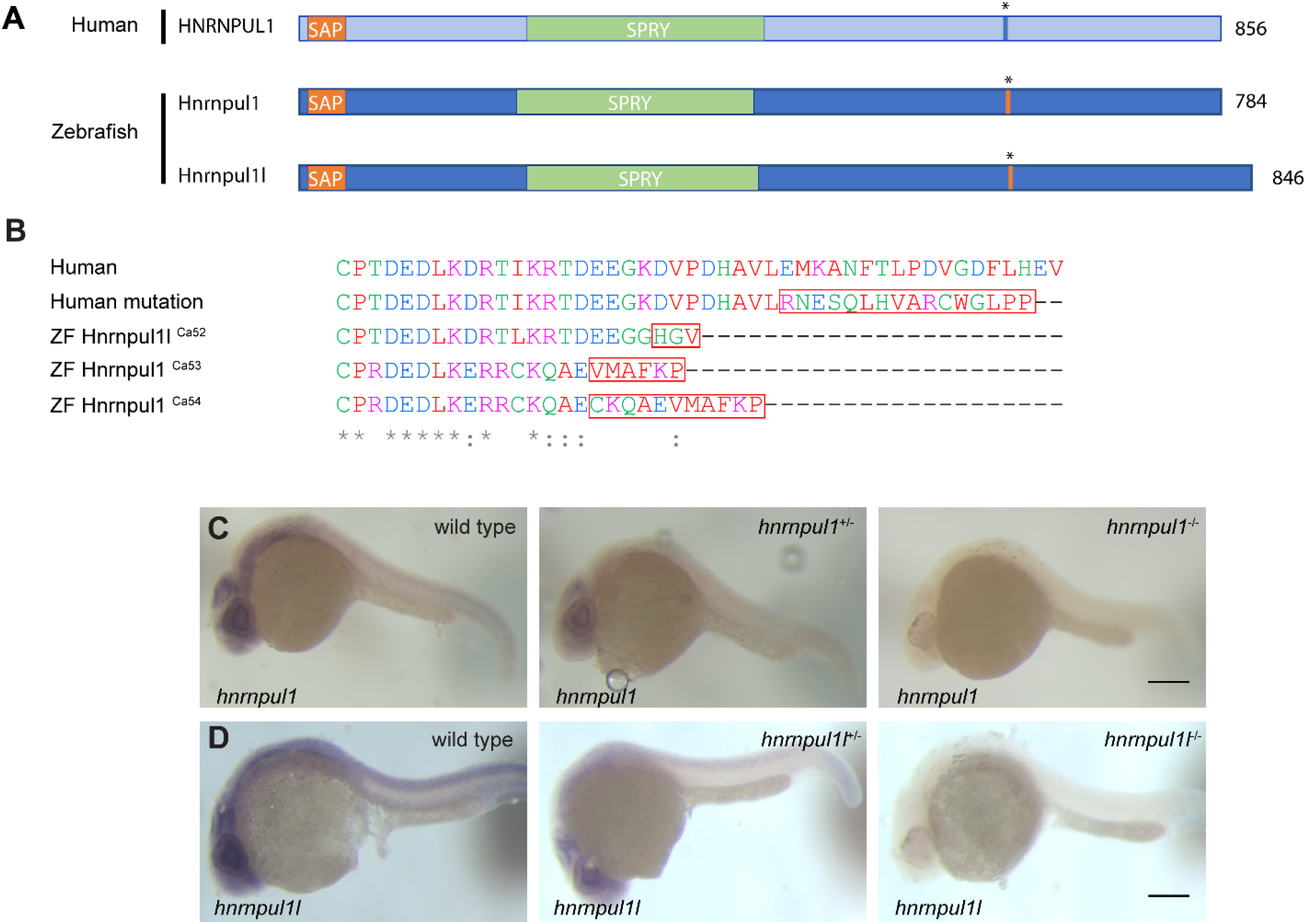
Mutation of human HNRNPUL1 and zebrafish hnrnpul1 and hnrnpul1l. A) Schematic showing domains of human HNRNPUL1 and zebrafish Hnrnpul1 and Hnrnpul1l proteins. The mutation location in human HNRNPUL1 and equivalent sequence in zebrafish is marked by *. B) Amino acid sequence of mutations, red boxes indicate nonsense sequence. C,D) whole mount in situ hybridisation (WISH) staining for hnrnpul1^Ca52^ (C) and hnrnpul1l^Ca53^ (D), at 24 hpf reveals expression that is not spatially restricted for both genes, and reduced expression due to nonsense mediated mRNA decay in mutants. Scale bars = 200 μm.

We determined the expression pattern of both *hnrnpul1* and *hnrnpul1l* in embryos at 24 hpf. Both transcripts are broadly expressed (Fig. 1C, D). *hnrnpul1* appears to have higher expression in the Central Nervous System (CNS). These results are in line with previous reports in the Zebrafish Information Network that expression is ‘not spatially restricted’ for *hnrnpul1* from 1 cell to Pec fin stage (60 hpf; Thisse and Thisse, 2004). No previous expression analysis is available for *hnrnpul1l*. The *hnrnpul1* and *hnrnpul1l* mutations result in nonsense-mediated decay of the transcript as whole mount *in situ* hybridisation (WISH) against *hnrnpul1* and *hnrnpul1l* in wild type, heterozygous and homozygous mutants (Fig. 1C,D), shows a decrease in mRNA expression, suggesting both are null alleles. Analysis of the gross morphology of *hnrnpul1/1l* double homozygous mutants showed low frequency developmental abnormalities including edema and embryo curvature (Fig. S2), however viable and fertile adults were obtained for all allelic combinations including *hnrnpul1/1l* double mutants.

### *hnrnpul1/1l* modulates alternative splicing and transcription

The hnRNP family regulates alternative splicing, but knowledge of the specificity and targets of *hnrnpul1* during development is limited. Thus, we performed paired-end bulk RNA sequencing (RNAseq) to identify differentially spliced events between wild type and *hnrnpul1/1l* mutant embryos at 3 dpf, a stage where fins and tendons are developing. We used VAST-TOOLS to identify splice junctions and characterise splicing events (Irimia *et al.*, 2014; Tapial *et al.*, 2017). We observed 76 alternative splicing (AS) events: 25 skipped exons, 33 retained introns, and 7 alternative 3 ‘splice site and 11 alternative 5’ splice site changes in mutants (Fig. 2, Table S1). The most differentially expressed exon in our AS analysis is exon 13 of *hnrnpul1l*, as expected from its excision via our CRISPR-Cas9 mutagenesis (Fig 2A, B).

**Figure 2.**
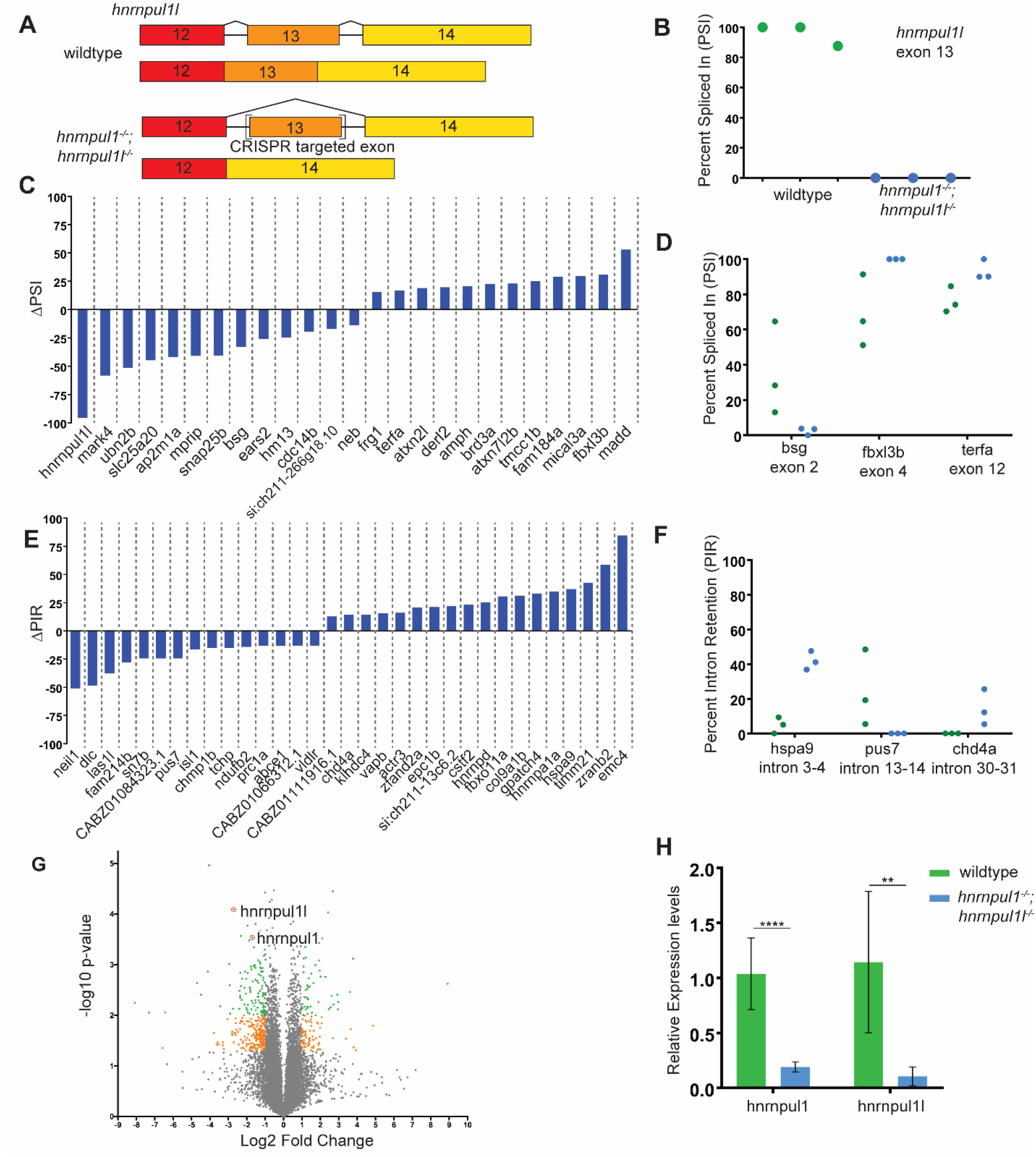
Loss of hnrnpu1l; hnrnpul1 leads to differential splicing. A) Schematic showing the processing of hnrnpul1l to form standard transcript and alternative splicing of exon 13 of the hnrnpul1l gene as a result of CRISPR-Cas9 targeted mutatgenesis. B) Percent Spliced In (PSI) for exon 13 of hnrnpul1l in wild type and hnrnpul1^−/−^; hnrnpul1l^−/−^ double mutant embryos at 3 dpf. C) Volcano plot showing all differentially expressed genes. Grey points = 1>Log2 FC>−1 or P>0.05, orange points = 1<Log2 FC<−1 and P≤0.05, green points = 1<Log2 FC<−1 and P≤0.01. D) qPCR validation of knock down of hnrnpul1 and hnrnpul1l expression in hnrnpul1^−/−^; hnrnpul1l^−/−^ double mutant embryos at 3 dpf compared to wild type. ** = P≤0.01, **** = P≤0.0001. E) Change in PSI of all exon skipping events in hnrnpul1^−/−^; hnrnpul1l^−/−^ double mutant embryos compared to wild type. F) Detailed view of PSI of genes associated with phenotypes, points represent each biological replicate. G) Change in Percent Intron Retention (PIR) of all intron retention events in hnrnpul1^−/−^; hnrnpul1l^−/−−^ double mutant embryos compared to wild type. H) Detailed view of PIR of genes associate with phenotypes, points represent each biological replicate. Details of affected exon/intron in Table S1.

Several genes with splicing changes and known developmental roles were identified (Fig 2C,E). *basigin (bsg)*, also known as CD 147, shows a 33% reduction in exon 2 usage in *hnrnpul1/1l* mutants compared to wild type. *F-box and leucine rich repeat protein 3 (fbxl3b)* shows a 31% increase in exon 4 usage in *hnrnpul1/1l* mutants compared to wild type, while *telomeric repeat binding factor a (terfa)* shows a 17% increase in exon 12 usage (Fig 2D). We also observed changes in Intron retention (Fig 2E). *Heat shock protein a 9 (hspa9)* shows a 37% increase in retention of intron 3-4, while *chromodomain helicase DNA binding protein 4a (chd4a)* shows a 14% increase in retention of intron 30–31 in *hnrnpul1/1l* mutants compared to wild type. *Pseudouridylate synthase 7 (pus7)* shows a 24% reduction in the retention of intron 13-14 in *hnrnpul1/1l* mutants compared to wild type (Fig. 2F).

Differential gene expression analysis identified 1575 genes that were significantly changed (p <0.05) with 1003 genes downregulated and 572 genes upregulated (Fig. 2G. Table S2). *hnrnpull1l* is downregulated 6.5 fold and *hnrnpul1* is downregulated 3 fold in this analysis consistent with our observations of nonsense-mediated decay (Fig 1C, D). To identify pathways that are dysregulated in in *hnrnpul1/1l* mutants we used Ingenuity Pathway Analysis (IPA). There is a significant upregulation of several pathways including translation (EIF4 and EIF2 signalling), protein ubiquitination, DNA damage via the 14-3-3 pathway, and the kinetochore metaphase pathways (Fig S3, Table S3). With respect to translation and ribosomal pathway changes, *eif1ax*, *eif33b* and *eif3k* factors are upregulated along with 14 *rpl* genes for the large ribosomal subunit and 6 *rps* genes for the small ribosomal subunit, and the 40S ribosomal protein S30, *fau* (Fig S3, Table S3). In the kinetochore-metaphase pathway, *cyclin B1*, cyclin dependent kinase *cdk1*, and the kinetochore protein *zwilch* are upregulated among other genes (Fig S3, Table S3). Although most pathways were upregulated (consistent with human HNRNPUL1 being a transcriptional repressor), we also found that early growth response genes *egr1* and *egr4* are downregulated 1.7- and 2.9-fold respectively, suggesting that cell division and differentiation might be reduced in mutants.

When we looked for dysregulated transcripts that may affect embryonic development, we found several key genes dysregulated. For instance, *glypican 6a* (*gpc6a*) expression is reduced 4-fold in *hnrnpul1/1l* mutants. *Gpc6a* is linked to omodysplasia in humans which includes shortening of extremities. We note reduced expression of *growth differentiation factor 3* (*gdf3*) an important growth factor in vertebral and skeletal development, downregulated 3.6-fold in in *hnrnpul1/1l* mutants. In terms of major developmental signalling pathways, notch pathway target *her15/hes5* is upregulated 4-fold, while *hedgehog acyltransferase-like (hhatlb)* is also upregulated 4-fold and *fgf11b* is downregulated 3-fold in mutants.

Thus, loss of hnrnpul1/1l leads to changes in both alternative splicing and in transcriptional regulation of genes involved in fundamental cellular processes (translation, ubiquitination and cell cycle) as well as key developmental patterning pathways.

### *hnrnpul1* mutation results in defects in craniofacial tendon development

We next assessed the developmental consequences of loss of *hnrnpul1/1l*. One of the most obvious phenotypes visible in mutant larvae is a gaping jaw at 8 dpf as compared wild types (Fig. 3A, A’). The incidence of an open jaw is significantly increased (wild type = 10%, *hnrnpul1^+/+^; hnrnpul1l^−/−^* = 29%, *hnrnpul1^+/−^; hnrnpul1l^−/−^* = 38%, *hnrnpul1^−/−^; hnrnpul1l^−/−^* =39%; Fig. 3B-D). We tested whether the gaping jaw is due to mispatterning of pharyngeal skeletal elements but found that Alcian blue staining shows cartilage is correctly patterned.

**Figure 3.**
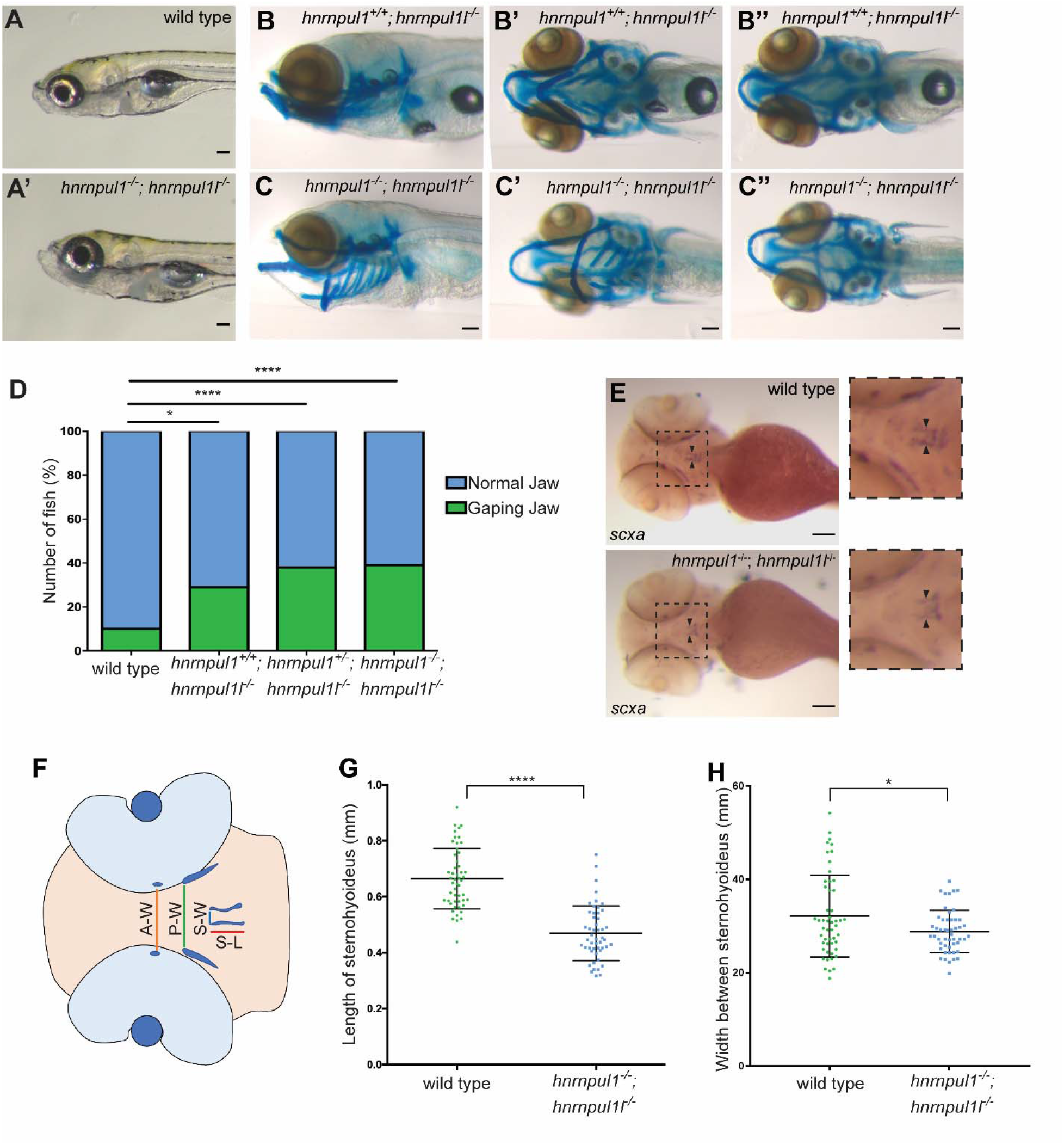
hnrnpu1/1l double mutants show a craniofacial phenotype due to shortened Sternohyoideus tendon. A) Images of live 8 dpf wild type and hnrnpul1^−/−^;hnrnpul1l^−/−^ double mutant larvae. B-C”) Alcian blue staining at 8 dpf. Lateral (B, C), ventral (B’, C’) and dorsal (B”, C”) views shown. Example of a normal jaw phenotype in hnrnpul1l^−/−^ single mutant (B) and a gaping jaw phenotype in hnrnpul1^−/−^;hnrnpul1l^−/−^ double mutant (C). D) Quantification of the proportion of fish showing a gaping jaw phenotype. Wild type n=283, hnrnpul1l^−/−^; hnRNPUL1^+/+^ n=24, hnrnpul1l^−/−^; hnrnpul1^+/−^ n=64, hnrnpul1^−/−^;hnrnpul1l^−/−^ n=84 from 5 trials. * = P≤0.05, **** = P≤0.0001, determined by Fisher’s test. E) WISH staining for scleraxis (scxa) in the Sternohyoideus tendon (arrow heads) in wild type and hnrnpul1^−/−^;hnrnpul1l^−/−^ double mutant embryos at 72 hpf. F) Schematic showing craniofacial tendons. Coloured lines demonstrate location of tendon measurements. A-W = width between Adductor Mandibulae tendons, P-W = width between Palatoquadrate tendons, S-W= width between Sternohyoideus tendons, S-L = Sternohyoideus length. G, H) Quantification of the length of, and width between the Sternohyoideus tendons in wild type (n= 51) and hnrnpul1^−/−^; hnrnpul1l^−/−^ double mutant (n= 50) embryos, from 3 trials. Scale bars = 100 μm. * = P≤0.05, **** = P≤0.0001 determined by Student’s T-test.

We next analysed the development of tendons in the embryonic craniofacial region that are responsible for opening and closing of the jaw (Fig 3F). Expression of *scleraxis* (*scxa*), a tendon specific marker at 72 hpf shows that *hnrnpul1/1l* mutants have a significantly shorter Sternohyoideus tendon (wild type = 0.66 ± 0.1 μm, *hnrnpul1/1l* = 0.47 ± 0.1 μm, p=<0.0001; Fig. 3E, G). The distance between the most anterior points of the Sternohyoideus tendons is also significantly narrower in *hnrnpul1/1l* mutants compared to wild types (wild type = 0.35 ± 0.09 μm, *hnrnpul1/1l* = 0.30 ± 0.05 μm, p=0.007; Fig. 3H). However, we find no difference in other craniofacial tendons, the Adductor Mandibulae or Palatoquadrate tendons (Fig. S4). These latter tendons are not responsible for jaw movement. Craniofacial tendons develop from neural crest cell (NCC) populations therefore we analysed the expression of the NCC markers *foxd3* and *sox10.* Expression of these specification markers at 12 hpf was normal in *hnrnpul1/1l* mutants (Fig. S5). Thus, the specification of craniofacial bone and cartilage appears normal, but failure of the Sternohyoideus tendon to grow properly may contribute to the gaping jaw phenotype by not allowing the mandible to close properly.

### *hnrnpul1* mutation results in increased incidence of scoliosis in adult zebrafish

We noted that young adult *hnrnpul1/1l* mutants show scoliosis. To determine whether this is congenital or idiopathic scoliosis we tested when scoliosis is first visible. 16 dpf larvae were stained with Alcian blue to visualise the maturing spinal column. Although *hnrnpul1/1l* mutant larvae are significantly smaller than wild type (Fig. S6), the development of the spinal column and vertebral structure is normal and comparable to wild type fish at this stage (Fig. 4A). We next compared the incidence of scoliosis in mutants and wild types at 16 weeks, when they are sexually differentiated young adults. We carefully controlled housing density, a factor that influences growth rates. Fish were housed at a density of 10 per 3L tank from 4.5 weeks until 16 weeks of age, at which point they were sacrificed and processed for alizarin red staining to visualise bone. The relative severity of spinal curvature was scored as none, mild, moderate, or severe scoliosis (Fig. 4B). The incidence of scoliosis is significantly higher in *hnrnpul1/1l* mutants (76%) compared to wild types (28%, p=<0.0001; Fig. 4C). There is no difference in the incidence of mild or moderate scoliosis between groups. However, the most severe phenotype only occurs in *hnrnpul1/1l* mutants.

**Figure 4.**
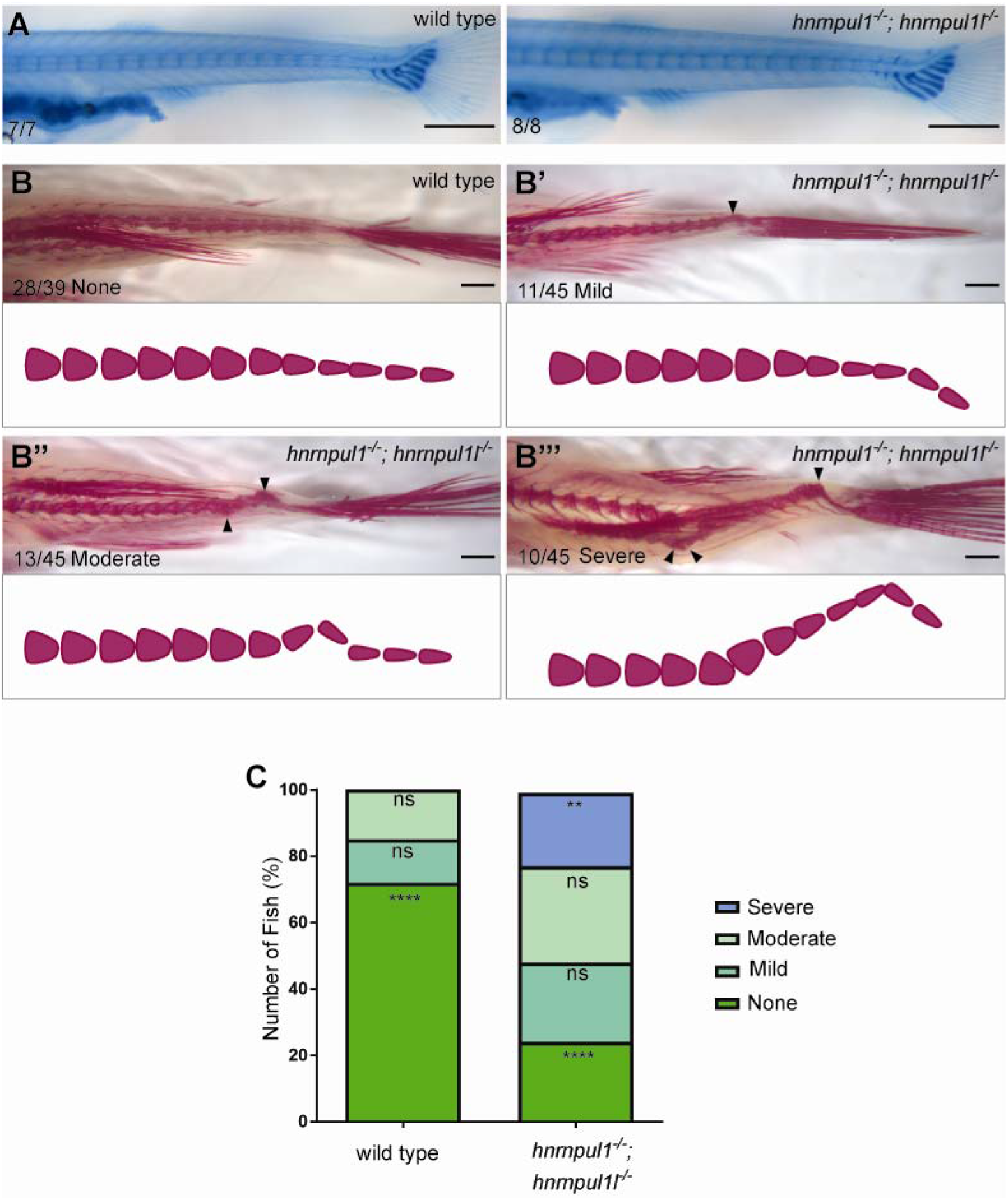
Loss of hnrnpu1l; hnrnpul1 leads to scoliosis. A) Alcian blue cartilage staining of wild type and hnrnpul1^−/−^; hnrnpul1l^−/−^ double mutant fish at 16 dpf showing normal larval spinal development. B-B’’’) Alizarin red bone staining of wild type (B) and hnrnpul1^−/−^; hnrnpul1l^−/−^ double mutant (B’-B’’’) fish at 16 weeks of age. Example images of relative mild (B’), moderate (B’’) and severe (B’’’) scoliosis with schematic to show curvature. C) Quantification of the proportion of fish with none, mild, moderate or severe scoliosis. Wild type n=18, hnrnpul1^−/−^; hnrnpul1l^−/−^ double mutant n=26, from 3 trials. ** = P≤0.01, **** = P≤0.0001 determined by Fisher’s test. Scale bar = 500 μm (A) 1000 μm (B-B’’’).

Overall, our data suggests that mutation of *hnrnpul1* and *hnrnpul1l* contributes to scoliosis that develops in the larval period, and is visible in the young adult. Because the phenotype is highly penetrant, it is likely congenital scoliosis.

### *hnrnpul1* mutation results in reduced fin growth but not fin specification

To understand whether *hnrnpul1* is involved in limb development, a common system affected by disrupted AS, we looked at the paralogous structure of paired fins of mutant zebrafish. We first determined if there is normal fin specification using expression of the limb specification markers *gli3, hand2* and *tbx5*. We found that expression of these genes is unchanged in mutants at 24-48 hpf suggesting that fins are specified normally (Fig. 5A-C’). We also tested expression of *wnt5b*, a wingless/wnt gene highly expressed in fin, but found that it was similarly unchanged in mutants (Fig 5D, D’). To determine if there were changes in embryonic fin size, we stained for *col1a1a,* which localises to the apical fold of the developing fin bud at 48 hpf (Fig. 5E, E’). *hnrnpul1/1l* mutants have significantly smaller fin bud area (8415 ± 3282 um^2^) compared to wild types (11350 ± 3338 um^2^, p=<0.0001; Fig. 5G). To ensure that decreased fin size was not due to defective overall growth we quantified eye size but find no significant difference between *hnrnpul1/1l* mutants and wild types, suggesting that the growth defect, at this stage, is specific to the fin and not the entire body (Fig. S7).

**Figure 5.**
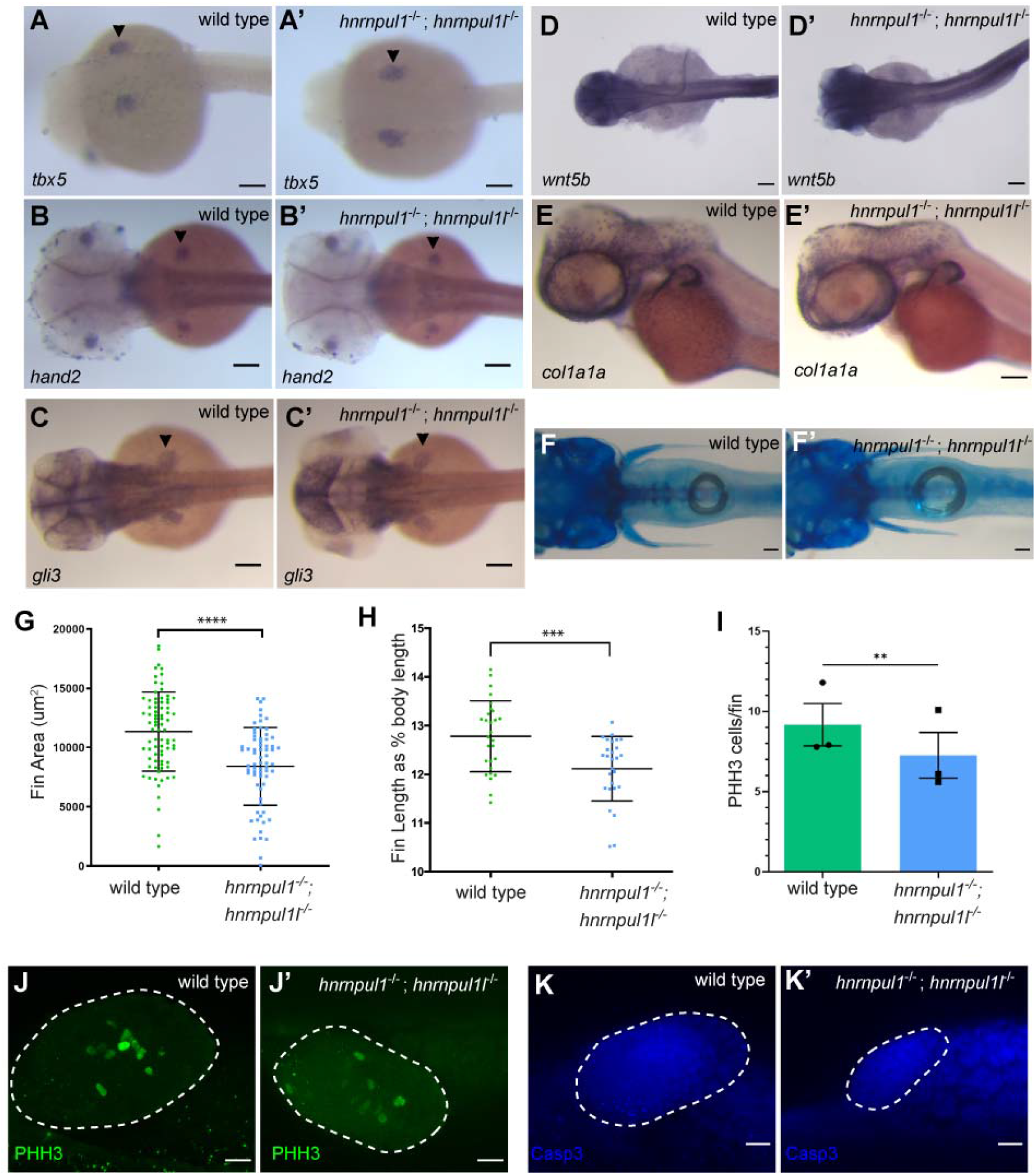
Loss of hnrnpul1 and hnrnpul1l does not affect fin specification, but leads to decreased fin growth in embryos and larvae. A-C’) mRNA expression of fin specification markers tbx5 (A, A’) at 24 hpf, hand2 (B, B’), gli3 (C, C’) and wnt5b (D, D’) at 48 hpf in wild type (A-D and hnrnpul1^−/−^;hnrnpul1l^−/−^ double mutant (A’-D’) embryos. D,D’). E) mRNA expression of col1a1a in wild type and hnrnpul1^−/−^;hnrnpul1l^−/−^ double mutant embryos at 49 hpf. F) Alcian blue cartilage staining of wild type and hnrnpul1^−/−^; hnrnpul1l^−/−^ double mutant fish at 16 dpf.E and F) Quantification of fin area in wild type and hnrnpul1^−/−^;hnrnpul1l^−/−^ double mutant col1a1a stained embryos at 49 hpf. Wild type n= 79, hnrnpul1^−/−^;hnrnpul1l^−/−^ n= 68, from 2 trials. H) Quantification of fin length at 16 dpf as a percentage of body length. Wild type n= 28, hnrnpul1^−/−^;hnrnpul1l^−/−^ n= 27, from 2 trials, determined using Student’s T-test. I) Quantification of proliferation via PhosphohistoneH3 (PHH3) immunostaining in wildtype and mutant fins at 48 hpf, N=3 experiments, n=30 fins, p=0.006 by the Paired Student’s T-Test. Dotted lines show the fin boundary. J-J’) Example PHH3 staining. K-K’) Example activated Caspase 3 (Casp3) immunostaining in wildtype and mutant fins. Scale bars = 100 μm. **=P ≤0.01, ***=P≤0.001, **** = P≤0.0001, ns = P>0.05.

We next tested whether fin growth is deficient at larval stages. We used Alcian blue staining at 16 dpf (Fig 5F, F’). Measuring pectoral fin length showed *hnrnpul1/1l* mutants have significantly shorter fins compared to wild types (Fig. S6). As larval fish differ in growth rates, to ensure smaller fin length was not due to overall smaller fish size we calculated the fin length as a proportion of standard length (tip to tail) for each animal. We find that *hnrnpul1/1l* mutants have significantly shorter fins compared to wild type with 12.8 ± 0.7% of body length in wild type animals and 12.1 ± 0.7% of body length in *hnrnpul1/1l* mutant animals (p=0.0008; Fig. 5H).

We assayed whether cell death of fin mesenchymal cells played a role in their smaller size by staining for activated Caspase 3 protein. There were no differences between wild type and mutant staining patterns (Fig 5K, K’). However, when we stained for a proliferation marker, phospho-Histone H3 (PHH3) which marks the nuclei of cells in mitosis, we detected a significant decrease from an average of 9 positive nuclei per fin at 48 hpf in wild type, to an average of 7 positive nuclei per fin in *hnrnpul1/1l* mutant (p= 0.006, Fig 5J, J’).

Taken together, our data suggest that *hnrnpul1* and *hnrnpul1l* play a role in regulating fin growth in embryos and larvae, but not in initial fin bud cell specification. Mechanistically, deficient proliferation in embryonic fins is one possible mechanism for the decreased fin size.

### *HNRNPUL1* as a candidate gene for human developmental anomalies

We found a variant of unknown significance (VUS) in HNRNPUL1 using whole exome sequencing of two similarly affected sisters with craniofacial and limb abnormalities. These sisters were born to consanguineous first cousin parents of Arab descent. Given the similarity between the two sisters but no family history of craniofacial or limb abnormalities, this suggested autosomal recessive or biallelic inheritance patterns. Our analysis identified the variant c.1673dup, p. (Glu560Argfs*17) in the gene *HNRNPUL1.* This gene has not been previously associated with a known genetic condition. We did not detect any additional patients with variants in this gene using web-based tools such as Matchmaker exchange (Buske *et al.*, 2015) or GeneMatcher, (Sobreira *et al.*, 2015).

The probands both presented at birth with multiple skeletal malformations following uncomplicated pregnancies. The older sister was born to a 27-year-old G4P4 (gravidity and parity) woman. Limb malformations were noted on a 27-week ultrasound. The child was born at 38 weeks gestation by spontaneous vaginal delivery with a birth weight of 3.8 kg (75-90^th^ percentile). Outside of limb malformations and dysmorphic features, the child’s neonatal examination and clinical course was unremarkable. She was born with bilateral short humeri, right absent ulna with two fixed in extension digits of the right hand; five rays present but with missing carpals, abnormal nails and dorsal creases on the left hand (Fig. 6A, B). Her lower limbs had mid shaft femoral pseudoarthroses, fused tibia to the femoral condyles, absent fibulas and abnormal toes (Fig. 6C, D). Her karyotype showed mosaic Turner syndrome, which is thought to be an incidental finding. Her course has been mostly uncomplicated except for orthopedic issues and intelligence is in the normal range. She is minimally dysmorphic with hypertelorism, upslanting palpebral fissures, prominent eyes and eyelashes, and a high palate.

**Figure 6.**
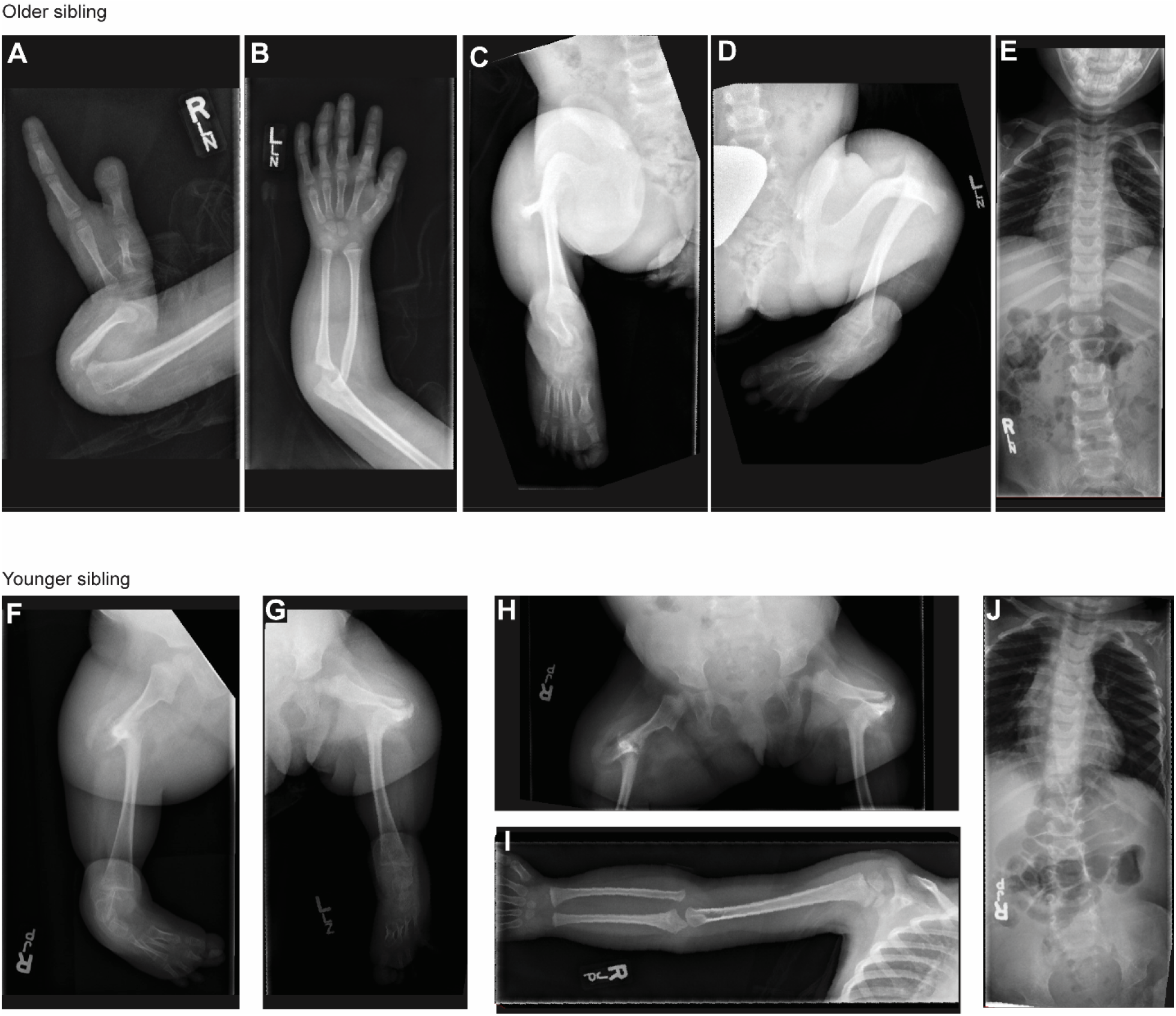
Radiographic features of siblings with a VUS in HNRNPUL1. A-E) Xray images of older sibling. Right arm showing short humerus and absent ulna with 2 fixed in extension digits of the right hand (A). Left arm showing short humerus and normal upper arm (B). Right (C) and Left (D) legs showing mid shaft femoral pseudoarthroses, fused tibia to the femoral condyles, absent fibulas and abnormal toes. F-J) X-ray images of the younger sibling Right (F) and Left (G) legs showing bilateral fibular agenesis, short and bowed femurs and four metarsals and tarsals (H). Right arm (I) showing normal upper limb development.

The younger sister was born at term to a 28-year-old G5P5 woman. Her birth weight was 4.03 kg (75-90^th^ percentile) with a head circumference of 39.0 cm (+2.5 SD). She had bilateral absent fibular, short and bowed femurs and four metatarsals and tarsals bilaterally (Fig. 6F, G, and H). Upper limb development was normal (Fig. 6I) with the exception of the hands showing stiff index finger dorsal creases and abnormal nails progressively more severe from ray 5 to 1, despite the presence of five rays. Her echocardiogram and abdominal ultrasounds were normal. She was felt to be minimally dysmorphic with a prominent forehead, relative macrocephaly, bitemporal narrowing, hypertelorism with prominent eyes and heavy eyelashes. Her palate was high but there was no clefting. Her clinical course has been mostly unremarkable outside of orthopedic complications and her intelligence is felt to be within the normal range. In addition to their skeletal malformations, both had bicornuate uterus, hirsutism, the first sister showed bilateral hydronephrosis, and the second sister showed Dandy Walker malformation. Clinically the girls share some features with Al-Awadi/Raas-Rothschild/Furhmann syndrome (OMIM 276820), which arises due to homozygous variants in *WNT7A* however *WNT7A* sequencing was negative.

Whole exome sequence analysis was undertaken as part of the Finding of Rare Disease Genes (FORGE) Canada consortium in both affected siblings. Variants were filtered based on rarity in the FORGE internal variant database and ExAC, along with predicted effects on protein function (Beaulieu *et al.*, 2014). Given the family structure, a recessive mode of inheritance was favoured, although heterozygous variants present in both siblings were assessed since germline mosaicism could not be excluded. Rare homozygous variants were identified in both affected siblings in the *PODXL, CFTR, HNRNPUL1, and PRKD2* genes. The variants PODXL, CFTR and PRKD2 were excluded as causative candidates (Table S3).

The rare homozygous variant in heterogeneous nuclear riboprotein U-like 1 (Chr19 (GRCh37):g.41807595dup, NM_007040.5 (HNRNPUL1):c.1673dup, p. (Glu560Argfs*17)) is predicted to result in a frameshift and premature stop approximately two-thirds of the way through the protein. No individuals with homozygous loss of function variants have been previously reported, and data from the Genome Aggregation Database (gnomAD) suggests that HNRNPUL1 loss of function variants are not tolerated in humans. Segregation of this variant by Sanger sequence analysis of both affected individuals, both parents, and four unaffected siblings shows that both patients are homozygous, both parents are heterozygous for the c.1673dup variant, while unaffected siblings are not homozygous for this variant.

## Discussion

Although we can detect changes in AS using sequencing technology, our understanding of how global mutations in splicing factors impact tissue-specific development and disease remains incomplete (Cieply and Carstens, 2015; Suzuki *et al.*, 2019). Phenotypes resulting from loss of *hnrnpul1/1l* in zebrafish are pleiotropic and affect multiple systems from craniofacial tendons to fin growth and skeletal morphology. This is not surprising as *hnrnpul1* and *hnrnpul1l* have a widespread expression pattern in embryonic zebrafish. It is particularly interesting that the phenotypes we observe are developmental, and specific to different organ systems rather than affecting the animal as a whole. We are the first to show that *hnrnpul1/1l* affects both alternative splicing (AS) and expression of mRNAs *in vivo*. Tissue specific phenotypes likely occur through varying composition of components of the spliceosome within different tissues (reviewed in (Baralle and Giudice, 2017)), and through varying target transcript expression in tissues.

We cannot verify that the human HNRNPUL1 VUS are pathogenic without identifying additional human variants and/or further studies with the patient mutations. However, the identification of human patients with HNRNPUL1 homozygous loss of function variants, and the viability of zebrafish *hnrnpul1/l* double mutants provides evidence that mutations in this gene are tolerated in animals but result in severe developmental consequences. Loss of this gene may be tolerated because, as a splicing regulator, even a complete loss of *hnrnpul1* function may not lead to complete loss or mis-splicing of targets. For instance, we show that there is a difference in the ratio but not a complete loss of splice variants in the zebrafish mutants. We use zebrafish *hnrnpul1/1l* mutants to shed light on the large number of pathways and genes regulated by this gene during development. Pleiotropic phenotypes and a large number of splicing and transcriptional changes downstream of *hnrnpul1* preclude us from identifying single genes as causative of the phenotypes we observe. However there are many potentially interesting genes and pathways identified by sequencing.

### Hnrnpul1 regulation of translation, growth, and cell cycle genes

Ingenuity pathway analysis identified the major pathways affected in Hnrnpul1 mutants as ‘EIF2 signaling’, ‘EIF4 and p706SK signaling’, ‘Kinetochore Metaphase signaling’, ‘Protein ubiquitination’, and ‘DNA Damage 13-3-3δ’. In all cases, multiple genes in these pathways were upregulated ~1,5- to ~2- fold in mutants at the RNA level. Our sequencing was conducted on whole animals, and thus there may be higher differential expression if we looked at single tissues.

In addition to upregulated ribosomal and growth pathways, we also observed some important genes for growth and morphogenesis that were downregulated 3-to 4-fold in mutants including *egr1* and *egr4*. At the same time, genes involved in polyubiquitination were also upregulated. The loss of *hnrnpul1* therefore leads to a complex set of changes in gene regulation that have opposing effects on growth. The upregulation of protein ubiquitination, however, and specifically proteins involved in polyubiquitination, suggests an upregulation of the proteosome and protein degradation. Similarly, changes in the relative proportions of ribosomal subunits can lead to translational dysregulation and reactive oxygen species production, leading to DNA damage (Kampen *et al.*, 2020). DNA damage proteins were also upregulated in our RNAseq analysis. Thus, our analysis paints a picture of dysregulated proteostasis in *hnrnpul1/1l* mutants, potentially leading to changes in other cellular functions.

We find increases in transcription of cyclins and cyclin dependent kinases. While this might result in increased cell division, this is not reflected in the *in vivo* phenotype, at least at the level of the fin where proliferation is decreased. It is possible that upregulated mRNAs are not all translated, or even if protein levels are increased, other pathways may compensate.

We find that members of the FGF, Nodal, Notch and Sonic Hedgehog signalling effectors (*fgf11, gdf3, her15/hes5, hhatl1*) are downregulated in *hnrnpul1* mutants. Gdf3 is a Nodal ligand important for mesodermal and endodermal tissues. Embryonic morphological phenotypes observed in *gdf3* mutants resemble the low frequency morphological defects we observe in *hnrnpul1* mutants (Bisgrove *et al.*, 2017; Pelliccia *et al.*, 2017). Her15 functions in the somite segmentation clock (Trofka *et al.*, 2012), and Hhat1 has developmental roles in the heart (Shi *et al.*, 2020). All published phenotypes of these genes have been in early development, and not at larval stages when new phenotypes may appear such as the craniofacial and scoliosis phenotypes that we observe.

### Hnrnpul1/1l modulates alternative splicing

Homology to other HNRNP genes suggested that HNRPUL1 might play a role in AS, but this has never been demonstrated directly. We provide the first evidence for its role in AS in zebrafish by demonstrating alternative exon usage, intron retention, and alternative 3’ and 5’ splice sites in *hnrnpul1/1l* mutants. We suggest that some of these might have phenotypic consequences. For instance, hnrnpul1*/1l* mutants are smaller at larval stages. Some of the genes we found that have alternative splicing changes in zebrafish have been associated with short stature in humans. *PUS7* (de Brouwer *et al.*, 2018), *CHD4A* (Weiss *et al.*, 2016), and *FBXL3* are associated with short stature in humans, and are orthologs of *pus7, chd4a* and *fbxl3a* identified in our study (Ansar *et al.*, 2019). The role of *hnrnpul1* in AS of *pus7* and *fbxl3a* should be investigated as possible causes of defective growth.

A large scale zebrafish mutagenesis screen identified *telemetric repeat factor a* (*terfa*) mutants as having a protruding jaw phenotype (Golling *et al.*, 2002). Interestingly, *terfa* has altered AS in *hnrnpul1/1l* mutants. While the function of *terfa* in normal jaw development is not understood, it is possible that disrupted splicing of this gene could be contributing to the phenotype in *hnrnpul1/1l* mutants.

Identification of altered AS of *chd4a* in *hnrnpul1/1l* mutants is particularly interesting. CHD4, a chromodomain helicase DNA binding protein also participates in DNA repair (Pan *et al.*, 2012). Loss of CHD4 impairs recruitment of BRCA1 to sites of DNA damage. This is intriguing because a second known role for human HNRNPUL1 is directly in DNA repair. HNRNPUL1 is recruited by the MRN complex to sites of DNA damage to promote DNA resection (Polo *et al.*, 2012), a role separate from its AS and RNA activity that we have examined here. Control of DNA repair directly at the double strand break site as well as through AS of essential DNA repair regulators such as CHD4 highlights a potential dual role for HNRNPUL1 in DNA repair. Whether DNA repair defects contribute to any of the phenotypes in embryonic or adult zebrafish *hnrnpul1/1l* mutants should be investigated, particularly as additional DNA repair components had altered gene expression in our RNAseq analysis at the mRNA level.

### Hnrnpul1 roles in craniofacial tendon development

*hnrnpul1/1l* mutants have an unusual craniofacial phenotype of an open jaw. Despite apparent displacement of skeletal elements, we observed no defects in skeletal development. We thus examined the development of tendons. Zebrafish craniofacial tendons share patterning and developmental pathways with mammalian tendons. Bone, cartilage and tendons of the face are derived from the neural crest. Developing tendons require expression of *scleraxis (scxa)* for specification, followed by expression of differentiation markers *tenomodulin (tnmd)* and *collagen (col1a)* (Chen and Galloway, 2014). Muscular and tendon development is coupled, as muscle attachment is required for the normal maturation of craniofacial tendons (Chen and Galloway, 2014) and mechanical load is needed for their differentiation (Brunt *et al.*, 2017). Zebrafish mutations leading to a disruption of craniofacial tendon differentiation have been linked to jaw closure defects (Mcgurk *et al.*, 2017). Zebrafish embryos that are anesthetised to prevent movement, show reduced jaw muscle activity and reduced tendon cell numbers in the jaw via reduced Wnt16 activity (Brunt *et al.*, 2017). We observe a shorter Sternohyoideus tendon, a tendon that connects the Sternohyoideus muscle to the hyohyal junction at the second pharyngeal arch midline (Mcgurk *et al.*, 2017). This region is referred to as the mandibulohyoid junction and is important for jaw opening. We find that tendons not associated with this junction (Adductor Mandibulae or Palatoquadrate tendons) show no difference in size, consistent with a specific defect in jaw tendon development at the midline as opposed to tendon development in general. A shorter Sternohyoideus tendon would hold the mandible open. Whether the Sternohyoideus is shorter due to decreased tenocyte specification and/or proliferation, or due to decreased activity and mechanical loading in *hnrnpul1/1l* mutants is unknown.

### Development of idiopathic scoliosis in *hnrnpul1* mutants

Scoliosis is defined as a three-dimensional rotation of the spine to an angle greater than 10°. Congenital scoliosis (CS) is present at birth arising due to a developmental abnormality, while idiopathic scoliosis (IS) develops during childhood or adolescence with no known cause (Goldstein and Waugh, 1973). The etiology of IS remains unknown but it does not appear to be due to vertebral abnormalities (Wajchenberg *et al.*, 2016). Neurological, muscular, growth and even hormonal abnormalities may be associated with IS (reviewed in (Latalski *et al.*, 2017)), however, no conclusive cause has been established. The fact that concordance is much higher in monozygotic twins than dizygotic twins does suggest a genetic link (Kesling and Reinker, 1997). Human IS has been difficult to study in mammalian models because common models such as mouse are quadrupeds and show a difference in spine structure and gravitational load (Ouellet and Odent, 2013). The zebrafish has recently emerged as an excellent model for IS, due to a similar cranial to caudal spinal load and the ease of genetic manipulation, which makes them susceptible to late onset spinal curvatures (Gorman and Breden, 2009). Therefore, we searched for a link between *hnrnpul1* and known genes leading to scoliosis in zebrafish. Zebrafish mutants in *ptk7*, a regulator of Wnt signalling required in ciliated lineages for cilia motility, show late onset scoliosis. Defective cerebrospinal fluid (CSF) flow as a result of *ptk7* mutation leads to scoliosis, potentially by inducing neuroinflammation (Hayes *et al.*, 2014; Grimes *et al.*, 2016; Van Gennip *et al.*, 2018). CSF circulation is also important for circulation of neurotensin neuropeptides (Zhang *et al.*, 2018) and mechanosensation by Pkd2l1 (Sternberg *et al.*, 2018); mutants in these genes develop IS. Thus, disruption in CSF circulation leads to multiple downstream consequences that can result in the failure of spine straightening. While we did not find changes in expression or splicing of *ptk7*, *urps* or *pkd2l1* using whole embryo RNA sequencing, it is possible that signal from tissue specific differences may have been masked by using bulk sequencing of whole embryo tissue. However, we do find decreased expression of *gdf3* in *hnrnpul1/1l* mutants. Mutations in human *GDF3* cause skeletal abnormalities including vertebral fusion and in some patients, mild scoliosis (Ye *et al.*, 2009). Additionally, as idiopathic scoliosis is late-developing, the RNAseq we undertook at 3 dpf may have been too early to detect changes relevant to motile cilia and CSF circulation. We did however find a decrease in expression of *arl3l2*. ARL3 proteins are present in the cilia and mutations lead to Joubert syndrome in humans (Alkanderi *et al.*, 2018; Powell *et al.*, 2019). Whether *arl3l2* plays a role in cilia activity in the zebrafish is currently unknown but may provide a mechanistic link between *hnrnpul1/1l* and scoliosis. The human patients are now entering adolescence and have not developed scoliosis, but we will continue to monitor them given that we noted an unexpected scoliosis phenotype in early adult *hnrnpul1*/*1l* zebrafish mutants.

### Involvement of Hnrnpul1 in fin and limb growth

Paired fins of teleosts (including zebrafish) are ancestral to limbs in tetrapods (including humans) as they are derived from locomotive organs in common ancestral vertebrates (Shubin *et al.*, 1997). Specification of limb and fin bud tissue is marked by expression of *tbx5* as early as 14-16 hours post fertilisation (hpf) in the zebrafish (Gibert *et al.*, 2006). Following specification, at approximately 23-26 hpf, condensation of mesenchymal cells forms the fin bud. Similar to mammals, the fish fin develops an apical ectodermal ridge (AER) as a signalling center driving mesenchymal cell proliferation and growth. In fish, the AER transforms into the apical fold at approximately 36 hpf. The limb/fin bud actively grows as mesenchymal cells organise and begin to differentiate into muscle masses around 46-48 hpf (Grandel and Schulte-Merker, 1998).

The roles of genes that initiate limb specification, such as retinoic acid, *tbx5*, and *fgf10* are conserved across vertebrates (Mercader, 2007). However, in zebrafish *hnrnupl1/1l* mutants there is no change in expression or splicing of limb specification genes. At this point we cannot conclude that the patients’ *HNRNPUL1* variant is the cause of their limb anomalies, as our analysis is correlative. However, there are some similarities to the zebrafish model. Siblings with the HNRNPUL1 VUS have shortened limbs, rather than an absence of limbs, consistent with *hnrnpul1* fish mutant data. The patients have variable loss and shortening of bones with some limbs unaffected and others severely affected. Overall, both patients show shortening of elements of the limb zeugopod (humerus and fibula) and variable agenesis of elements in the stylopod (ulna and tibia). The fish does not have correlates of these bones, however mesenchymal outgrowth that forms the fins and limbs (including bones) occurs via a homologous process. Mechanistically, we show a decrease in cell proliferation, but no change in cell death, in *hnrnpul1/1l* mutant fins at the same developmental stage where we observe smaller fins. This observation suggests that a reduction in normal proliferation and cell cycling prevents fins from growing at a normal rate.

Growth of the limb occurs via proliferation of limb mesenchyme, driven by FGF signals from the Apical Ectodermal Ridge (AER) in fish and tetrapods. Fgf10 induces Fgf8 via Wnt3 in chick, mouse and zebrafish (Mercader, 2007; Yano and Tamura, 2013). Although both fish and other vertebrates rely on Fgf signalling, the expression pattern of specific FGFs in different species is slightly different (reviewed in (Yano and Tamura, 2013)). We used RNA sequencing to identify transcripts and splice variants targeted by *hnrnpul1/1l* that might affect limb growth and identified some interesting candidates. We identified differential alternative splicing between *hnrnpul1/1l* mutant and wild type fish in *basigin* (*bsg*; also known as CD147) exon 2. Bsg exon 2 encodes a 351 bp/ 117 amino acid immunoglobulin domain, one of 3 Ig domains present in this transmembrane glycoprotein. Exon 2 is also alternatively spliced in human Bsg (Karczewski *et al.*, 2019). Bsg/CD147 is stabilised by CBRN, a chaperone that can be bound by thalidomide, a known teratogen that reduces limb size (Eichner *et al.*, 2016). Knockdown of *bsg/cd147* in zebrafish reduces pectoral fin size and phenocopies the teratogenic effects of thalidomide, suggesting that reduction in bsg expression impairs fin growth. Thalidomide is known to have anti-proliferative effects, while *bsg/CD147* promotes proliferation and invasiveness of cancer cells *in vitro* (Yang *et al.*, 2017). The effect of AS of *bsg* on fin/limb development has not been explored in any species.

The three strongest phenotypes in the zebrafish mutants (fin defects, craniofacial tendon development and scoliosis) all have links to Wnt signalling, even though RNA sequencing did not detect any defects in genes in these pathways. The patients with the HNRNPUL1 VUS appear to have a very similar phenotype to Fuhrmann or Al-Awadi/Raas-Rothschild/Schinzel phocomelia syndromes, both caused by loss of WNT7A (Woods *et al.*, 2006). Furthermore, pathogenic variants in WNT5A in humans lead to autosomal dominant Robinow syndrome, and mutations in its receptor ROR2 lead to recessive Robinow syndrome in humans (Van Bokhoven *et al.*, 2000; Person *et al.*, 2010). Robinow syndrome is characterised by craniofacial defects, short stature and vertebral segmentation defects. Although we have not detected members of the canonical Wnt pathway undergoing AS in *hnrnpul1* mutants, the phenotypes that we observe are very consistent with disruption of Wnt signalling. It is possible that Wnt is regulated secondarily through other *hnrnpul1* targets, potentially including cilia genes such as *arl3l2*. Thus, even though the link is between *hnrnpul1* and Wnt signaling is still unknown, the phenotypic similarities and affected developmental structures suggest similar underlying mechanisms should be explored.

## Materials and Methods

### Animal and patient data

All Zebrafish (*Danio rerio*) strains were maintained and raised under established protocols (Westerfield, 2000) and all animal procedures were approved by the University of Calgary Animal Care Committee (protocol AC17-0189). Zebrafish embryos were collected and incubated at 28.5°C in E3 medium (5 mM NaCl, 170 μM KCl, 330 μM CaCl_2_, 300 μM MgSO_4_, 223μM Methylene blue) and staged in hours post fertilisation (hpf) or days post fertilisation (dpf). When required, endogenous pigmentation was inhibited from 24 hpf by addition of 0.003% 1-phenyl-2-thiourea (PTU, Sigma Aldrich) in E3 medium.

This study was part of the Finding of Rare Disease Genes in Canada (FORGE Canada) consortium and approved by the University of Calgary Conjoint Health Research Ethics Board (REB# 23927).

DNA was extracted from all family members from whole blood using Puregene chemistry (Qiagen). Exome capture was undertaken in both affected individuals using the SureSelect 50 Mb All Exon Kit v3 (Agilent) followed by sequencing with a HiSeq2000 (Illumina). Variant calling and annotation were as described in Lynch et al. (2014). Confirmation and segregation of the *HNRNPUL1* c.1406dup variant was performed by PCR amplification (HotStar Taq Plus, Qiagen, Toronto, ON) from genomic DNA and Sanger sequencing with the ABI BigDye Terminator Cycle Sequencing Kit v1.1 (Life Technologies, Burlington, ON) on a 3130xl genetic analyzer (Life Technologies). Sequence subtraction and analysis was performed using Mutation Surveyor software (SoftGenetics, State College, PA).

### Generation of *hnrnpul1* and *hnrnpul1l* mutant Zebrafish

CRISPR mutations were created in both the *hnrnpul1* and *hnrnpul1l* genes by injection of guide RNAs at the single cell stage in conjunction with Cas9 mRNA and a homology-directed repair STOP cassette sequence oligonucleotide (Table S4). Guide RNAs were designed to target a location close to the human mutation. Founders were identified by genomic PCR analysis using primers in Table S4. DNA from F1 heterozygotes was cloned into pCR™Blunt II-TOPO^®^ vector (Thermofisher) and sequenced. Mutants are genotyped by PCR using primers described in Table S4, detailed protocol in Supplementary methods.

### Whole mount *in situ* hybridisation (WISH)

Embryos were fixed in 4% paraformaldehyde in PBS with 0.1% Tween-20 (PFA) at 4°C overnight and stored in 100% methanol at −20°C until required. All whole-mount *in situ* hybridisation was carried out according to standard protocols (Lauter *et al.*, 2011). Antisense probes for *hnrnpul1*, *hnrnpul1l*, *scxa*, *hand2, tbx5, foxd3* and *sox10* were produced by in vitro transcription using T7 polymerase (Roche), in the presence of digoxigenin-11-UTP (Sigma), from PCR fragments amplified from embryonic zebrafish cDNA (Table S4). Antisense probes for *gli3* and *col1a1a* were a gift from Peng Huang and produced from plasmid clones. Images were taken on a Zeiss Stemi SV 11 microscope with a Zeiss Axiocam HRc camera. Area and length measurements were completed in ImageJ using the line and measure tools or Zen Blue (Zeiss) using the line tool. Confocal images were collected on a Zeiss LSM700 confocal microscope and assembled in ImageJ and Adobe Photoshop.

### Alcian blue and Alizarin red staining

Alcian blue staining of 16 dpf zebrafish was carried out as previously described (Walker and Kimmel, 2007). In brief, fish were fixed in 4% PFA overnight at 4°C and stained in 0.04% Alcian blue in 100 mM Tris-HCl/ 10 mM MgCl_2_. Following staining, fish were washed in decreasing concentration of Ethanol/100 mM Tris-HCl to remove excess stain. Fish were bleached in 3% H_2_0_2_/ 0.5% KOH until pigment was lost. Fish were then washed in increasing concentrations of glycerol in 1% KOH until 100% glycerol. Fish were imaged in 100% glycerol.

Alizarin red staining of adult zebrafish was carried out as previously described (Connolly and Yelick, 2010). In brief, adult zebrafish were eviscerated and fixed for 48hrs in 4% PFA at 4°C. Zebrafish were bleached in 30% H_2_0_2_/ 1% KOH for 2 hours followed by 2 hours in 15% H_2_0_2_/ 0.5% KOH. Zebrafish were cleared in 1% trypsin/2% borax solution overnight and stained in 1mg/ml alizarin red in 1% KOH over-night. Following staining fish were washed in increasing concentrations of glycerol in 1% KOH until 100% glycerol. Fish were imaged in 100% glycerol on a Zeiss Stemi SV11 microscope.

### Antibody staining

Embryos were fixed in 4% PFA in PBT overnight and placed in methanol overnight at −20°C. They were permeabilized with acetone (Sigma) at −20°C for 20 minutes before transfer to PBT for 3× 5 minutes, then blocked in PBT plus 5% normal sheep serum (Sigma) for 1 hour. Primary antibodies (Phospho-histone H3 (Ser10) from EMD Millipore Corporation 06570, or active Caspase 3 from BD Pharmingen 559565) were diluted 1/250 in blocking solution and applied overnight. After washing for 2 hours in multiple changes of PBT, Goat anti Rabbit-Alexa 488 secondary antibody (Thermo Fisher, A-11008) was applied for 2 hours in blocking solution. Embryos were washed with multiple changes of PBT overnight before being visualized using a Zeiss LSM700 confocal microscope. Cells positive for PHH3 were counted blinded.

### RNA sequencing

Zebrafish were genotyped from excised tail tissue, while matching head tissue from the same embryo was snap frozen at −80°C for RNA extraction after genotyping. Tails were exposed to 25mM NaOH at 55°C for 30 minutes then neutralised with 40mM Tris HCl pH5 to extract genomic DNA (Meeker *et al.*, 2007) followed by PCR genotyping. Eight embryos of each genotype were pooled per replicate and total RNA was purified using RNeasy Plus mini kit (Qiagen). Three replicates each of wild type sibling and *hnrnpul1/1l* mutants were sequenced using paired end reads on Illumina NextSeq500 to a read depth of ~100M reads. RNA libraries were prepared using NEBNext Ultra II Directional RNA Library Prep kit (New England Biolabs). Alternative splicing analysis of RNA sequencing data was completed using the Vertebrate Alternative Splicing and Transcript Tools (VAST-TOOLS) v2.2.2, (Irimia *et al.*, 2014; Tapial *et al.*, 2017). For all events, a minimum read coverage of 10 actual reads per sample was required, as described (Irimia *et al.*, 2014) using genome release danRer10 (Torres-Méndez *et al.*, 2019). PSI values for single replicates were quantified for all types of alternative events, including single and complex exon skipping events (S, C1, C2, C3, ANN), microexons (MIC), alternative 5’ss and 3’ss (Alt5, Alt3) and retained introns (IR-S, IR-C). A minimum ΔPSI of 10% was required to define differentially spliced events upon each knockdown, as well as a minimum range of 5% between the PSI values of the two samples.

Differential gene expression analysis was performed using RPKM output from VAST-TOOL analysis. For each gene p-values were determined by Student’s T-test of RPKM values from three biological replicates. Log_2_ FC was calculated using the mean RPKM for each genotype. A p-value of ≤0.05 were required to define a gene as significantly differentially expressed. Volcano plots and heat maps were performed using GraphPad Prism. Gene Ontology and Pathway analysis used Qiagen Ingenuity Pathway Analsys.

### Statistical analysis

All experiments were performed in at least three independent biological replicates. All quantitative data are presented as mean ± standard deviation. Statistical analysis was performed using PRISM Graph Pad Software. ns, P>0.05, *P≤0.05, **P≤0.01, ***P≤0.001, ****P≤0.0001.

## Supporting information

Blackwell Supplement

## Acknowledgements

We would like to thank the patients and their family for participation in this research. We thank the Alberta Children’s Hospital Research Institute for funding for DB and SF through the MORPH project. Operating funding was received from the Canadian Institute of Health Research Institute of Genetics Rare Disease Models and Mechanisms (180309-001-001) to SJC.

Patient sequencing was performed under the Care4Rare Canada Consortium funded by Genome Canada, the Canadian Institutes of Health Research, the Ontario Genomics Institute, Ontario Research Fund, Genome Alberta, Genome BC, Genome Quebec, and Children’s Hospital of Eastern Ontario Foundation.

We would also like to thank the MORPH committee for guidance, and Juan Valcárcel Juárez and Wendy Dean for helpful comments on the manuscript.

## Data availability

All RNASeq reads have been deposited with the National Centre for Biotechnology Information in the Gene Expression Omnibus database and are available under the accession GSE144754.

