## Supplementary material for "Hnrnpul1 controls transcription, splicing, and modulates skeletal and limb development in vivo": Blackwell Supplement

**A**

```
Human               GACGAGGAAGGGAAGGATGTCCCAGATCATGCGGT-CTTAGAAATGAAAGCCAACTTCAC
Human_Mutation      GACGAGGAAGGGAAGGATGTCCCAGATCATGCGGTTCTTAGAAATGAAAGCCAACTTCAC
                    *********************************** ************************

Human               GTTGCCAGATGTTGGGGACTTCCTCCATGAGGTT
Human_Mutation      GTTGCCAGATGTTGGGGACTTCCTCCATGAGGTT
                    **********************************
```

**B**

```
ZF_hnrnpul1l        GATGAGGAAG--------------------------------------------------
ZF_hnrnpul1l_Ca52   GATGAGGAAGGTCATGGCGTTTAAACCTTAATTAAGCTTAATTAATTAATTAATTAATTA
                    **********

ZF_hnrnpul1l        -------------------------------------------------------GGAAG
ZF_hnrnpul1l_Ca52   ATTAAGCTTAATTAATTAAGCTGTTGTAGGGAAGGATGTACAACAGCTGTTGTAGGGAAG
                                                                           *****

ZF_hnrnpul1l        GATGTTCCCGATCATGCTGTATTAGAAATGAAAG
ZF_hnrnpul1l_Ca52   GATGTTCCCGATCATGCTGTATTAGAAATGAAAG
                    **********************************
```

**C**

```
ZF_hnrnpul1         GTAAGCAAGCTGAG-----------------------------------GATGGGAAAGA
ZF_hnrnpul1_Ca53    GTAAGCAAGCTGAGGTCATGGCGTTTAAACCTTAATTAAGCTGTTGTAAGATGGGAAAGA
                    **************                                   ***********

ZF_hnrnpul1         TGTGCCCGATCAAGCCGTTTTAGAAATGAAAG
ZF_hnrnpul1_Ca53    TGTGCCCGATCAAGCCGTTTTAGAAATGAAAG
                    ********************************
```

**D**

```
ZF_hnrnpul1         GTAAGCAAGCTGAG-----------------------------------------------
ZF_hnrnpul1_Ca54    GTAAGCAAGCTGAGTGTAAGCAAGCTGAGGTCATGGCGTTTAAACCTTAATTAAGCTGTTG
                    **************

ZF_hnrnpul1         ----------------GATGGGAAAGATGTGCCCGA-TCAAGCCGTTTTAGAAATGAAAG
ZF_hnrnpul1_Ca54    TAGATGGGAAAGATGTGATGGGAAAGATGTGCCCGATTCAAGCCGTTTTAGAAATGAAAG
                                    ******************** ***********************
```

**Figure S1 – *hnrnpul1* and *hnrnpul1l* wild type and mutation DNA alignments**

A) Human wild type and mutated *HNRNPUL1*. B) Zebrafish wild type and CRISPR-Cas9 mutated allele Ca52 of *hnrnpul1l*. C, D) Zebrafish wild type and CRISR-Cas9 mutated allele Ca53 (C) and Ca54 (D) of *hnrnpul1*. Yellow highlights location of duplicated T seen in patient mutation. Red highlights premature STOP resulting from the mutation. Green box highlights insertion due to CRISPR-Cas9 mutagenesis.

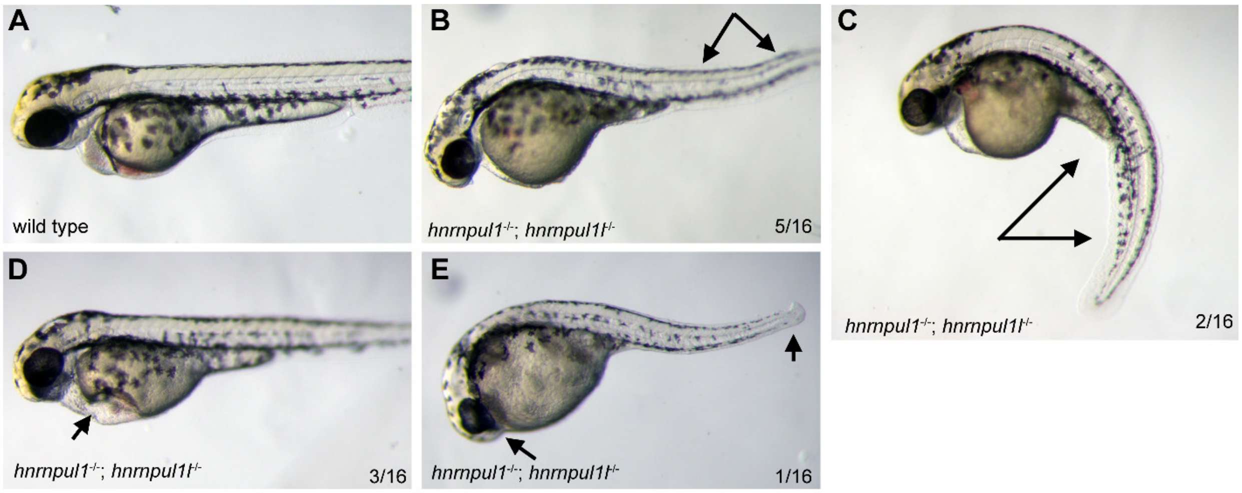


**Figure S2– Low frequency severe developmental phenotypes in *hnrnpul1/1l* mutants at 48hpf**

Gross morphology of wild type (A) and *hnrnpul1*$^{-/-}$;*hnrnpul1l*$^{-/-}$ double mutants at 48hpf (B-E). While the majority of embryos are viable and survived to adulthood, a minority of embryos show developmental phenotypes including B) Dorsal curvature C) Ventral curvature D) Edema E) Missing heart and caudal fin.

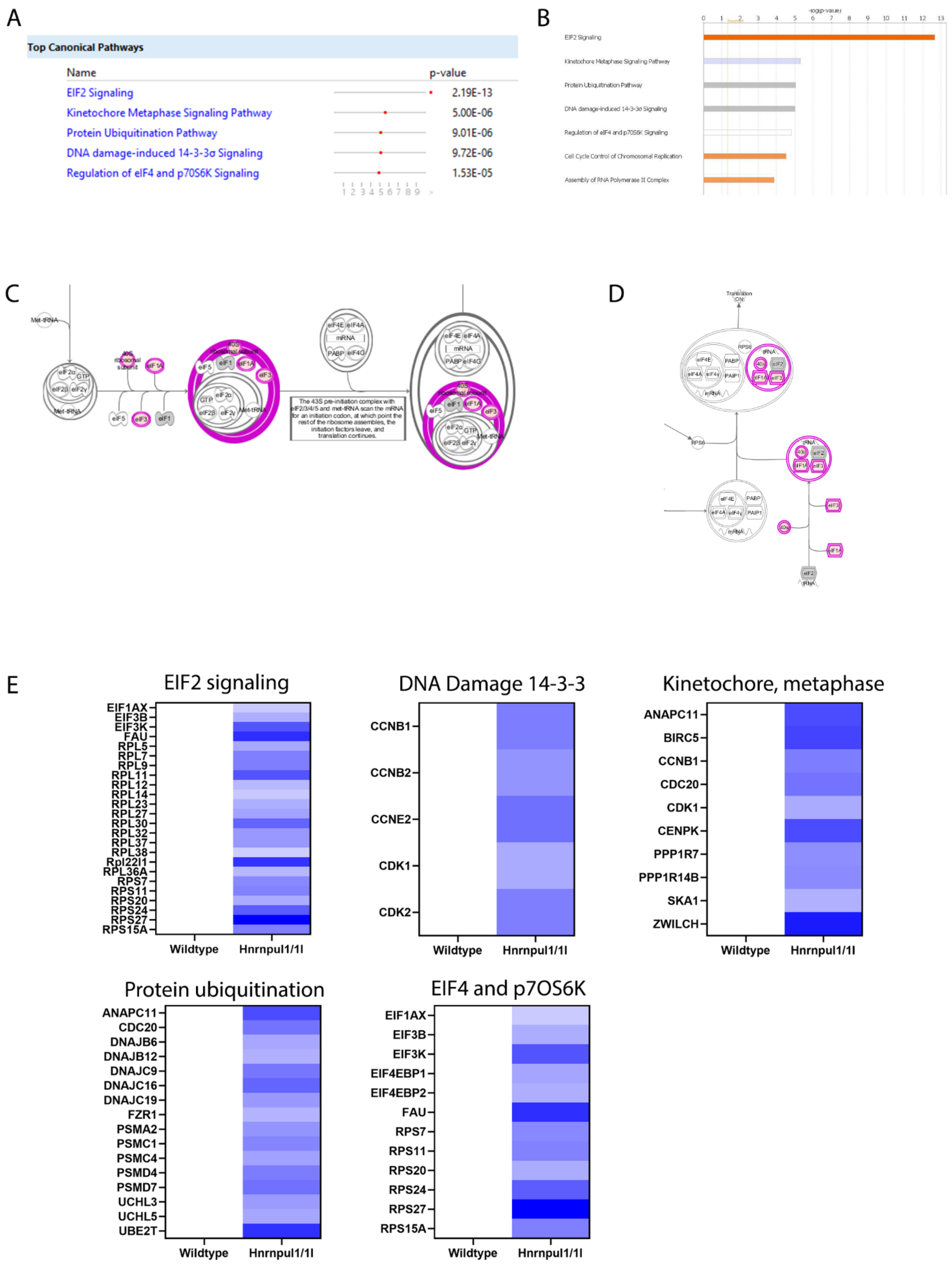
A
Top Canonical Pathways
Name
p-value
EIF2 Signaling 2.19E-13
Kinetochore Metaphase Signaling Pathway 5.00E-06
Protein Ubiquitination Pathway 9.01E-06
DNA damage-induced 14-3-3σ Signaling 9.72E-06
Regulation of eIF4 and p70S6K Signaling 1.53E-05
B
-log(p-value)
0 1 2 3 4 5 6 7 8 9 10 11 12 13
EIF2 Signaling
Kinetochore Metaphase Signaling Pathway
Protein Ubiquitination Pathway
DNA damage-induced 14-3-3σ Signaling
Regulation of eIF4 and p70S6K Signaling
Cell Cycle Control of Chromosomal Replication
Assembly of RNA Polymerase II Complex
C
Met-tRNA
40S ribosomal subunit
The 43S pre-initiation complex with eIF2/3/4/5 and met-tRNA scan the mRNA for an initiation codon, at which point the rest of the ribosome assembles, the initiation factors leave, and translation continues.
D
Translation ON
E
EIF2 signaling
EIF1AX
EIF3B
EIF3K
FAU
RPL5
RPL7
RPL9
RPL11
RPL12
RPL14
RPL23
RPL27
RPL30
RPL32
RPL37
RPL38
Rpl22l1
RPL36A
RPS7
RPS11
RPS20
RPS24
RPS27
RPS15A
Wildtype
Hnrnpul1/1l
DNA Damage 14-3-3
CCNB1
CCNB2
CCNE2
CDK1
CDK2
Wildtype
Hnrnpul1/1l
Kinetochore, metaphase
ANAPC11
BIRC5
CCNB1
CDC20
CDK1
CENPK
PPP1R7
PPP1R14B
SKA1
ZWILCH
Wildtype
Hnrnpul1/1l
Protein ubiquitination
ANAPC11
CDC20
DNAJB6
DNAJB12
DNAJC9
DNAJC16
DNAJC19
FZR1
PSMA2
PSMC1
PSMC4
PSMD4
PSMD7
UCHL3
UCHL5
UBE2T
Wildtype
Hnrnpul1/1l
EIF4 and p7OS6K
EIF1AX
EIF3B
EIF3K
EIF4EBP1
EIF4EBP2
FAU
RPS7
RPS11
RPS20
RPS24
RPS27
RPS15A
Wildtype
Hnrnpul1/1l

**Figure S3–*hnrnpul1/1l* mutants show expression changes in pathways controlling fundamental cell processes by Ingenuity Pathway Analysis** A) The top 5 pathways identified as disrupted in *hnrnpul1/1l* mutants. B) graphical representation of most highly changed pathways. C) Purple highlights the steps in EIF2 translational processes disrupted in *hnrnpul1/1l* mutants. D) Purple highlights disruptions EIF2 and p70S6K pathway in *hnrnpul1/1l* mutants. E) Heatmaps showing upregulation of genes in the marked pathways (Baseline in wild type is white, and increased expression over baseline is color coded in increasingly dark blue with 2 fold being darkest blue).

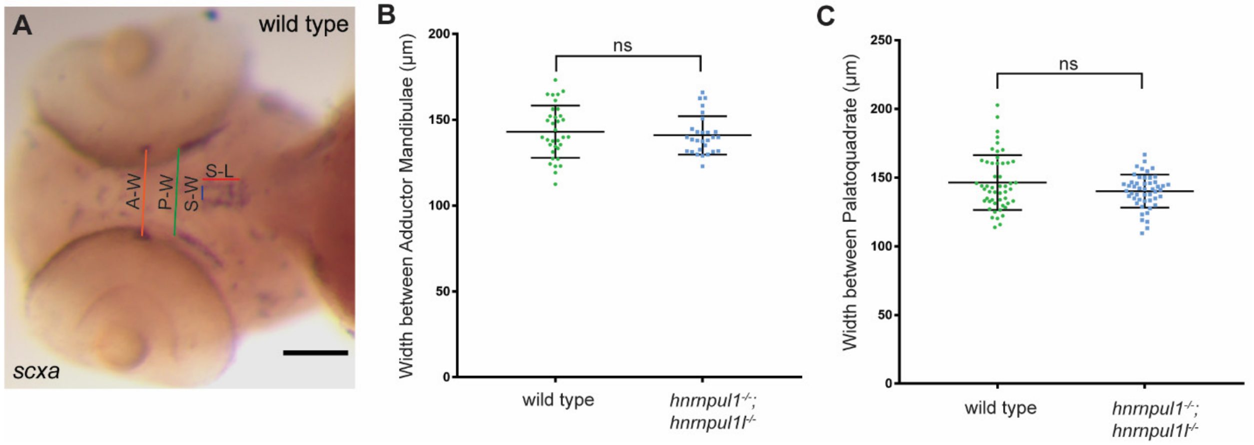


**Figure S4 – *hnrnpu1/1l* mutants do not show differences in the width between the Palatoquadrate or Adductor Mandibulae tendons.**

A) WISH staining for *scleraxis* (*scxa*) in the Sternohyoideus, Palatoquadrate and Adductor Mandibulae tendons in a wild type embryo at 72hpf. Coloured lines demonstrate location of tendon measurements. A-W = width between Adductor Mandibulae tendons, P-W = width between Palatoquadrate tendons, S-W= width between Sternohyoideus tendons, S-L = Sternohyoideus length. B) Quantification of the width between Adductor Mandibulae tendons in wild type (n=34) and *hnrnpul1*$^{-/-}$; *hnrnpul1l*$^{-/-}$ double mutant (n=28) embryos. C) Quantification of the width between Palatoquatrade tendons in wild type (n=51) and *hnrnpul1*$^{-/-}$; *hnrnpul1l*$^{-/-}$ double mutant (n=49) embryos. ns = not significant, determined by Student T-test.

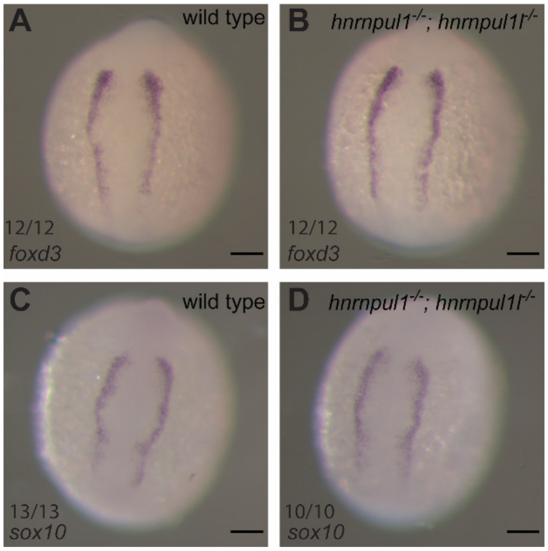


**Figure S5 – Early neural crest cell specification is not affected by *hnrpul1* mutations**

A,B) WISH staining for *foxd3* in wildtype (A) and *hnrnpul1*$^{-/-}$; *hnrnpul1l*$^{-/-}$ double mutant (B) embryos at 12hpf. C,D) WISH staining for *sox10* in wildtype (C) and *hnrnpul1*$^{-/-}$; *hnrnpul1l*$^{-/-}$ double mutant (D) embryos at 12hpf. Scale bars = 100µm.

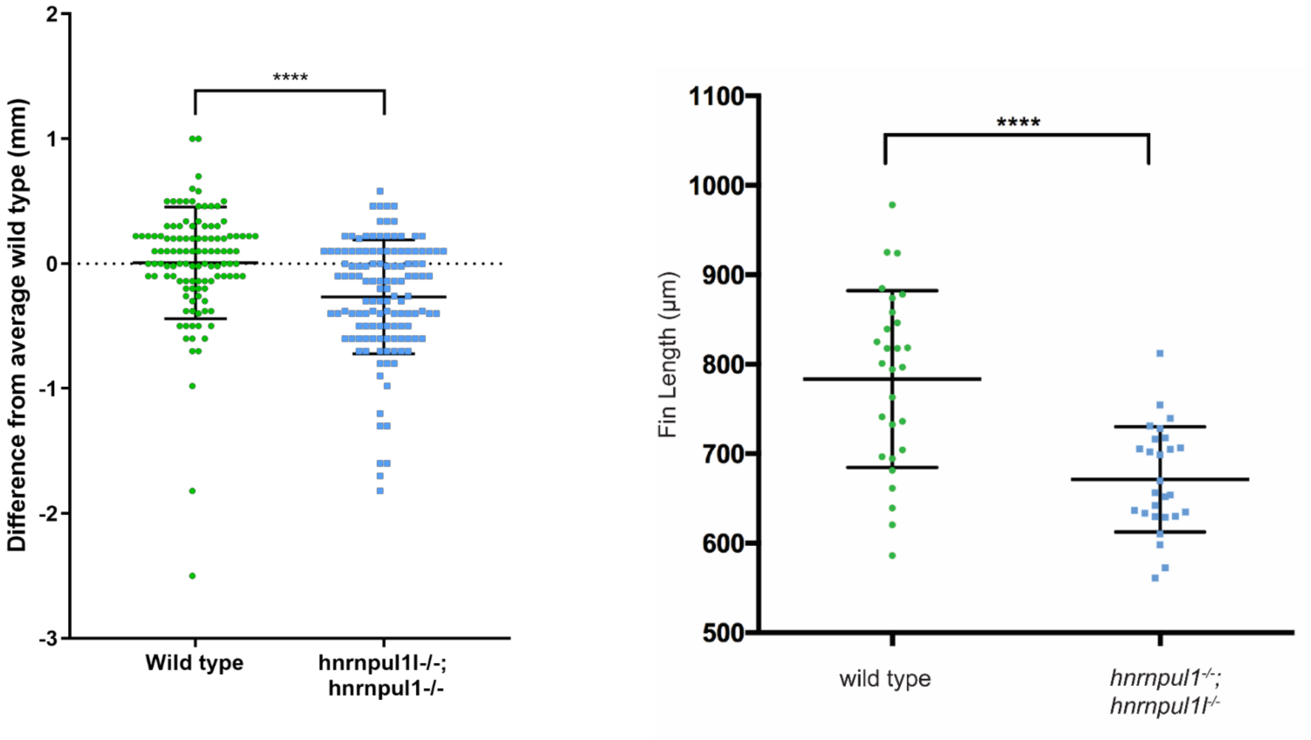


**Figure S6 - *hnrnpu1/1l* double mutants are smaller and have smaller pectoral fins at larval stages compared to wild types.**

Quantification of standard length normalised to the mean wild type standard length of 16dpf larvae. Wild type n= 115, *hnrnpul1*$^{-/-}$; *hnrnpul1l*$^{-/-}$ double mutant n= 127. **** = P<0.0001, determined by Student's T-test. Quantification of pectoral fin length at 16dpf .Wild type n= 28, *hnrnpul1*$^{-/-}$;*hnrnpul1l*$^{-/-}$ n= 27. **** = P<0.0001, determined by Student's T-test.

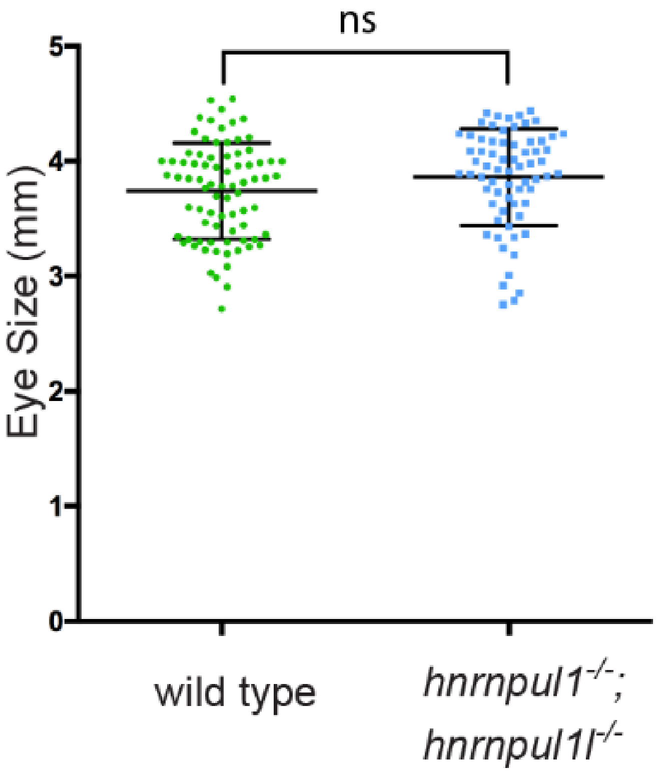


**Figure S7: Eye size is not changed in hnrnpul1/1l mutants despite body growth defects.**

Eye size was measured from the indicated genotypes from 16 dpf larvae in the longest dimension. There is no statistical difference in eye size between mutant and wildtype.

**Table S3 – Non-causative variants identified in patients with limb anomalies.**

| Gene | Variant | Reason for exclusion |
|---|---|---|
| Podocalyxin-like gene *POXDL* | Chr7(GRCh37):g.131195974C>T, NM_001018111.2(PODXL): c.319G>A, p.(Val107Met) | • A biallelic loss of function variant in this gene was reported in an autosomal recessive juvenile Parkinson family (1)<br>• Homozygous knockout in mouse results in perinatal lethality due to severe defects in kidney development and omphalocele. Limb development is normal in these embryos (2) |
| protein kinase D2 gene *PRKD2* | Chr19(GRCh37):g.47204104G>A, NM_016457.4(PRKD2): c.1073C>T, p.(Ala358Val) | • *in silico* analysis (SIFT, PolyPhen2, alignGVGD, MutationTaster) did not predict any damaging effect on the protein and either heterozygous or homozygous loss of PRKD2 in Rhesus monkey and mouse, respectively, leads to hyperinsulinemia and insulin resistance without any reported congenital limb anomalies (3). |
| cystic fibrosis transmembrane conductance regulator gene *CFTR* | Chr7(GRCh37):g.117232214A>T, NM_000492.3(CFTR): c.1993A>T, p.(Thr665Ser) | • This variant has been reported as disease-associated in the literature where is has been found in the heterozygous state in two clinically affected individuals of Tunisian and Egyptian descent (4,5)<br>• *In vitro* functional studies suggest potentially decreased chloride currents (6) and a splice enhancer effects leading to partial exclusion of coding sequence (7).<br>• This finding was unexpected as neither affected individual presented with clinical features of cystic fibrosis, although this variant may be associated with a mild presentation. Given the paucity of clinical information on the impact of this variant in the literature, it is impossible to determine if homozygosity for this variant has any clinical impact. Nevertheless given the well studied nature of pathogenic variation in *CFTR* in humans, this gene is not a plausible candidate for the striking developmental anomalies in these siblings. |

**Table S4 – Primers used for CRISPR mutagenesis, genotyping, production of WISH probes and qPCR.**

| Gene Name | Forward Primer | Reverse Primer | Product Size | Use |
|---|---|---|---|---|
| *hnrnpul1l* | TAATACGACTCACTATAGCGAACTGATGAGGAAGGGAGTTTTAGAGCTAGAAATAGCAAG | N/A | N/A | sgRNA |
| *hnrnpul1l* | AAAGCGAACTGATGAGGAAGGTCATGGCGTTTAAACCTTAATTAAGCTGTTGTAGGGAAGGATGTTCCCGATCAT | N/A | N/A | STOP cassette |
| *hnrnpul1* | TAATACGACTCACTATAGGTGTAAGCAAGCTGAGGATGTTTTAGAGCTAGAAATAGCAAG | N/A | N/A | sgRNA |
| *hnrnpul1* | GAAGGTGTAAGCAAGCTGAGGTCATGGCGTTTAAACCTTAATTAAGCTGTTGTAGGATGGGAAAGATGTGCCCGA | N/A | N/A | STOP cassette |
| *hnrnpul1l* | AGGTTTTCAACGCAAGGCTA | GGGTGGGTTCTTCCAAGTCT | WT= 238bp,<br>Ca52= 344bp | Genotyping |
| *hnrnpul1* | GGTTTCACCGTAAGGCTGTT | TTTGTAGCATCCCATTTTTCA | WT= 246 bp<br>Ca53= 281bp<br>Ca54= 309bp | Genotyping |
| *hnrnpul1* | AACAGCCCACCTGTGATGAG | GCTCACTGCTGGGAAGATGT | 672bp | WISH |
| *hnrnpul1l* | TCTTCCAGGAGCAGAAAAAGCA | GCACCTGCCCACAAATAAGC | 538bp | WISH |
| *scleraxis* | ACAGAGACAGAAAGCCGGAGGAGT | CTTACCATTTTCCTCTGGTTGCTGAG | 850bp | WISH |
| *hand2* | TCGCTGTCATGAAGAACCCC | GGCCAACCAGTTCTCCCTTT | 513bp | WISH |
| *tbx5* | AAACTCTCCCAGTGACAGCG | TGTGTGTTCGTGGTAGGAGC | 913bp | WISH |
| *col1a1a* | AAGGAGGGCCAGAAAGGTAA | AGGGTGGTGTCAACCTCAAG | 995bp | WISH |
| *gli3* | (8) | | | WISH |
| *sox10* | ACCGTGACACACTCTACCAAGATGACC | TAATACGACTCACTATAGGCATGATAAAATTTGCACCCTGAAAAGG | 935bp | WISH |

| | | | | |
|---|---|---|---|---|
| *foxd3* | CGGCATTGGGAATCCATA | TAATACGACTCACTATAGGCAACGAAATGAAATAGAAAGAAGGA | 684bp | WISH |
| *wnt5b* | GGATTACTGCCTGCGCAATG | **TGTAATACGACTCACTATAC**AGCTCTGACATCAGCAAGGT | 659 bp | WISH |
| *hnrnpul1* | AGACGTCAGCTTGGAGAACA | CAGCATGTTTAGCAGCCCAT | 125bp | qPCR |
| *hnrnpul1l* | AAATCACGGCAGATCCCAGA | CTTCACATCATAACGCCCGG | 75bp | qPCR |

**Supplementary methods:**

### Genotyping *hnrnpul1/1l* mutants

Genomic DNA was prepared by exposing tissue to 25mM NaOH at 55°C for 30 mins (50µl for adult fin clips and whole embryos, 25µl for embryo tails). The solution was neutralised with an equal volume of 40 mM Tris HCl pH=5. Samples were vortexed and centrifuged, stored at -20°C until required for PCR. Final PCR reaction consisted of 2 µl 5X Phusion HF buffer (New England Biolabs, Massachusetts – B0518S), 0.2 µl 10 mM dNTPs (ThermoFisher), 0.5 µl 50 µM forward primer, 0.5 µl 50 µM reverse primer (Table S1), 0.1 µl Phusion polymerase (NEB – M0530), 5.7 µl $H_2O$ and 1 µl gDNA. PCR reactions were subject to the following conditions 98°C for 30 secs; 98°C for 10 secs, 62°C for 20 secs, 72°C for 30 secs (35 cycles); 72°C for 3 mins. PCR products were visualised on 2% agarose electrophoresis gel, expected product sizes are detailed in Table S3.
